# Dynamic nuclear structure emerges from chromatin crosslinks and motors

**DOI:** 10.1101/2020.08.22.262758

**Authors:** Kuang Liu, Alison E. Patteson, Edward J. Banigan, J. M. Schwarz

## Abstract

The cell nucleus houses the chromosomes, which are linked to a soft shell of lamin filaments. Experiments indicate that correlated chromosome dynamics and nuclear shape fluctuations arise from motor activity. To identify the physical mechanisms, we develop a model of an active, crosslinked Rouse chain bound to a polymeric shell. System-sized correlated motions occur but require both motor activity *and* crosslinks. Contractile motors, in particular, enhance chromosome dynamics by driving anomalous density fluctuations. Nuclear shape fluctuations depend on motor strength, crosslinking, and chromosome-lamina binding. Therefore, complex chromatin dynamics and nuclear shape emerge from a minimal, active chromosome-lamina system.

---

The cell nucleus houses the genome, or the material containing instructions for building the proteins that a cell needs to function. This material is ~ 1 meter of DNA with proteins, forming chromatin, and it is packaged across multiple spatial scales to fit inside a ~ 10 *μ*m nucleus [1]. Chromatin is highly dynamic; for instance, correlated motion of micron-scale genomic regions over timescales of tens of seconds has been observed in mammalian cell nuclei [2–6]. This correlated motion diminishes both in the absence of ATP and by inhibition of the transcription motor RNA polymerase II, suggesting that motor activity plays a key role [2, 3]. These dynamics occur within the confinement of the cell nucleus, which is enclosed by a double membrane and 10-30-nm thick filamentous layer of lamin intermediate filaments, the lamina [7–9]. Chromatin and the lamina interact through various proteins [10–12] and form structures such as lamina-associated domains (LADs) [13, 14]. Given the complex spatiotemporal properties of a cell nucleus, how do correlated chromatin dynamics emerge and what is their interplay with nuclear shape?

Numerical studies suggest several explanations for correlated chromatin motions. A confined Rouse chain with long-range hydrodynamic interactions that is driven by extensile dipolar motors can exhibit correlated motion over long length and timescales [4]. Correlations arise due to the emergence of local nematic ordering of within the confined globule. However, such local nematic ordering has yet to be observed. In the absence of activity, a confined heteropolymer may exhibit correlated motion, with anomalous diffusion of small loci [15, 16]. However, in marked contrast with experimental results [2, 3], introducing activity in such a model does not alter the correlation length at short timescales and decreases it at longer timescales.

Since there are linkages between chromatin and the lamina, chromatin dynamics may influence the shape of the nuclear lamina. Experiments have begun to investigate this notion by measuring nuclear shape fluctuations [17]. Depletion of ATP, the fuel for many molecular motors, diminishes the magnitude of the shape fluctuations, as does the inhibition of RNA polymerase II transcription activity by *α*-amanitin [17]. Other studies have found that depleting linkages between chromatin and the nuclear lamina, or membrane, results in more deformable nuclei [18, 19], enhanced curvature fluctuations [20], and/or abnormal nuclear shapes [21]. Interestingly, depletion of lamin A in several human cell lines leads to increased diffusion of chromatin, suggesting that chromatin dynamics is also affected by linkages to the lamina [22]. Together, these experiments demonstrate the critical role of chromatin and its interplay with the nuclear lamina in determining nuclear structure.

To understand these results mechanistically, we construct a chromatin-lamina system with the chromatin modeled as an *active* Rouse chain and the lamina as an elastic, polymeric shell with linkages between the chain and the shell. Unlike previous chain and shell models [20, 23, 24], our model has motor activity. We implement the simplest type of motor, namely extensile and contractile monopoles, representative of the scalar events addressed in an earlier two-fluid model of chromatin [25]. We also include chromatin crosslinks, which may be a consequence of motors forming droplets [26] and/or complexes [27], as well as chromatin binding by proteins, such as heterochromatin protein I (HP1) [28]. Recent rheological measurements of the nucleus support the notion of chromatin crosslinks [23, 24], as does indirect evidence from chromosome conformation capture (Hi-C) [29]. In addition, we explore how the nuclear shape and chromatin dynamics mutually affect each other by comparing results for an elastic, polymeric shell with those of a stiff, undeformable one.

## Model

Interphase chromatin is modeled as a Rouse chain consisting of 5000 monomers with radius *r*_*c*_ connected by Hookean springs with spring constant *K*. We include excluded volume interactions with a repulsive, soft-core potential between any two monomers, *ij*, and a distance between their centers denoted as 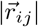, as given by 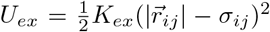 for 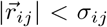, where 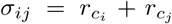, and zero otherwise. We include *N*_*C*_ crosslinks between chromatin monomers by introducing a spring between different parts of the chain with the same spring constant as along the chain. In addition to (passive) thermal fluctuations, we also allow for explicit motor activity along the chain. In simulations with motors, we assign some number, *N*_*m*_, of chain monomers to be active. An active monomer has motor strength *M* and exerts force **F**_***a***_ on monomers within a fixed range. Such a force may be attractive or “contractile,” drawing in chain monomers, or alternatively, repulsive or “extensile,” pushing them away (Fig. 1). Since motors *in vivo* are dynamic, turning off after some characteristic time, we include a turnover timescale for the motor monomers *τ*_*m*_, after which a motor moves to another position on the chromatin.

**FIG. 1.**
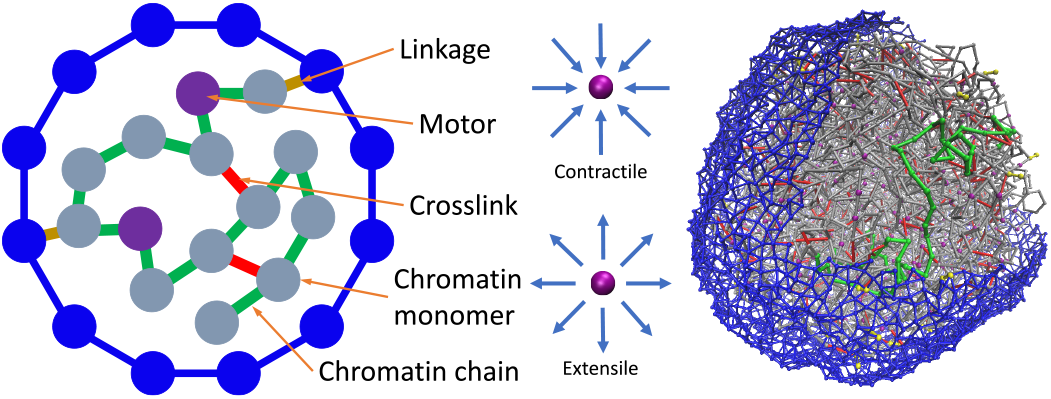
Left: Two-dimensional schematic of the model. Center: Schematic of the two types of motors. Right: Simulation snapshot.

The lamina is modeled as a layer of 5000 identical monomers connected by springs with the same radii and spring constants as the chain monomers and an average coordination number *z* ≈ 4.5, as supported by previous modeling [20, 23, 24] and imaging experiments [7–9]. Shell monomers also have a repulsive soft core. We model the chromatin-lamina linkages as *N*_*L*_ permanent springs with stiffness *K* between shell monomers and chain monomers (Fig. 1).

The system evolves via Brownian dynamics, obeying the overdamped equation of motion: 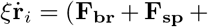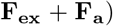, where **F**_***br***_ denotes the (Brownian) thermal force, **F**_***sp***_ denotes the harmonic forces due to chain springs, chromatin crosslink springs, and chromatin-lamina linkage springs, and **F**_***ex***_ denotes the force due to excluded volume. We use Euler updating, a time step of *dτ* = 10^−4^, and a total simulation time of *τ* = 500. For the passive system, **F**_***a***_ = 0. In addition to the deformable shell, we also simulate a hard shell by freezing out the motion of the shell monomers. To assess the structural properties in steady state, we measure both the radial globule, *R*_*g*_, of the chain and the self-contact probability. After these measures do not appreciably change with time, we consider the system to be in steady state. See SM for these measurements, simulation parameters, and other simulation details.

## Results

We first look for correlated chromatin motion in both hard shell and deformable shell systems. We do so by quantifying the correlations between the displacement fields at two different points in time. Specifically, we compute the normalized spatial autocorrelation function defined as 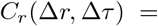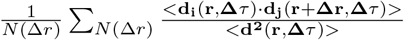, where Δ*τ* is the time window, Δ*r* is the distance between the centers of the two chain monomers at the beginning of the time window, *N* (Δ*r*) is the number of *ij* pairs of monomers within distance Δ*r* of each other at the beginning of the time window, and **d**_**i**_ is the displacement of the *i*^th^ chain monomer during the time window, defined with respect to the origin of the system. Two chain monomers moving in the same direction are positively correlated, while monomers moving in opposite directions are negatively correlated.

Fig. 2 shows *C*_*r*_(Δ*r*, Δ*τ*) for passive and active samples in both hard shell (Figs. 2 (a) and (b)) and soft shell cases for *N*_*C*_ = 2500, *N_L_* = 50, and *M* = 5 (Figs. 2 (e) and (f)). Both the passive and active samples exhibit short-range correlated motion when the time window is small, *i.e.*, Δ*τ* < 5. However, for longer time windows, both the extensile and contractile active samples exhibit more long-range correlated motion than the passive case. These correlations are visible in quasi-2d spatial maps of instantaneous chromatin velocities, which show large regions of coordinated motion in the active, soft shell case (Figs. 2 (c) and (g)).

**FIG. 2.**
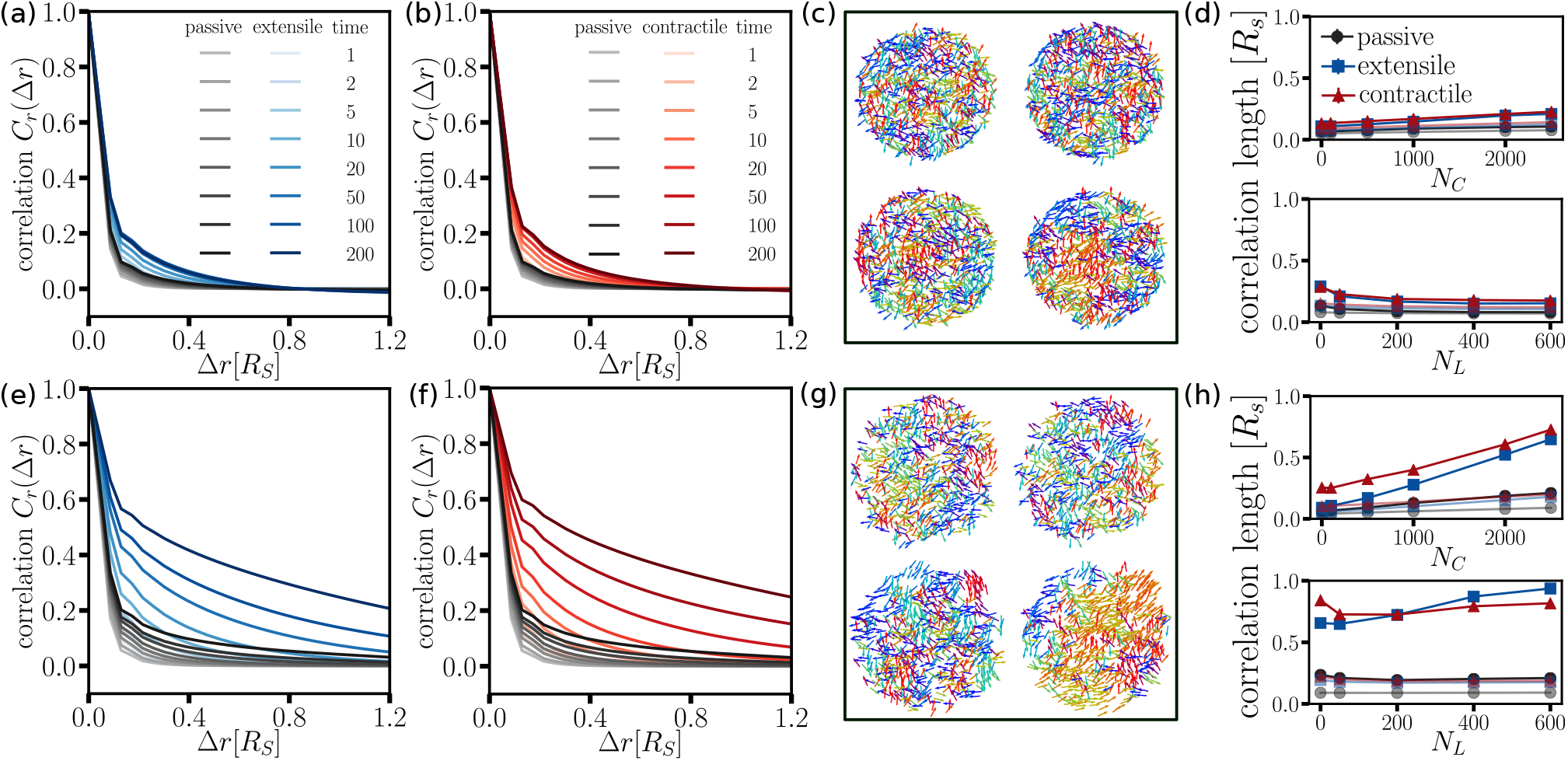
(a) The spatial autocorrelation function *C*_*r*_ (Δ*r*, Δ*τ*) for passive and extensile cases at different time lags, Δ*τ*, for the hard shell, while (b) shows the contractile and passive case. (c) Two-dimensional vector fields for Δ*τ* = 5 (left), 50 (right) for the passive case (top) and the contractile case (bottom). (d) The correlation length as a function of *N*_*L*_ and *N*_*C*_ for the two time lags in (c). (e~h): The bottom row shows the same as the top row, but with a soft shell. See SM for representative fits to obtain the correlation length.

To extract a correlation length to study the correlations as a function of both *N*_*C*_ and *N*_*L*_, we use a Whittel-Marten (WM) model fitting function 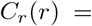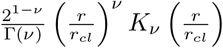 for each time window (Fig. 2 (f)) [3]. The parameter *ν* is approximately 0.2 for all cases studied. For the hard shell, the correlation length decreases with number of linkages (Fig. 2 (d)). This trend is opposite in deformable shell case with activity and long time lags (Fig. 2 (h)). For the hard shell, linkages effectively break up the chain into uncorrelated regions. For the soft shell, the shell deforms in response to active fluctuations in the chain. For both types of shells, the correlation length increases with the number of crosslinks (Figs. 2 (d) and (h)), with a more significant increase in the soft shell active case. It is also interesting to note that the lengthscale for the contractile case is typically larger than that of the extensile case, at least for smaller numbers of linkages.

Given the differences in correlation lengths between the hard and soft shell systems, we looked for net motion of the system in the soft shell case. Net motion has been observed in active particle systems confined by a deformable shell [30]. Similarly, we observe the active chain system moving faster than diffusively (see SM). In the shell’s center-of-mass frame, the correlation length is decreased, but still larger than in the hard shell simulations (see SM). Interestingly, experiments demonstrating large-scale correlated motion measure chromatin motion with an Eulerian specification (*e.g.*, by particle image velocimetry) and do not subtract off the global center of mass [2, 3, 6]. However, one experiment noted that they observed drift of the nucleus on a frame-to-frame basis, but considered it negligible over the relevant time scales [3]. Additionally, global rotations, which we have not considered, could yield large-scale correlations.

We also study the mean-squared displacement of the chromatin chain to determine if the experimental feature of anomalous diffusion is present. Figs. 3 (a) and (c) show the mean-squared displacement of the chain with *N*_*L*_ = 50 and *N*_*C*_ = 2500 as measured with reference to the center-of-mass of the shell for both the hard shell and soft shell cases, respectively. For the hard shell, the passive chain initially moves subdiffusively with an exponent of *α* ≈ 0.5, which is consistent with an uncrosslinked Rouse chain with excluded volume interactions [31]. However, the passive system crosses over to potentially glassy behavior after a few tens of simulation time units. We present *N*_*C*_ = 0 case in the inset to Fig. 3 for comparison to demonstrate that crosslinks are potentially driving a gel-sol transition as observed in prior experiments [32]. The active hard shell samples exhibit larger displacements than passive samples, with *α* ~ 0.6 initially before crossing over to a smaller exponent at longer times.

**FIG. 3.**
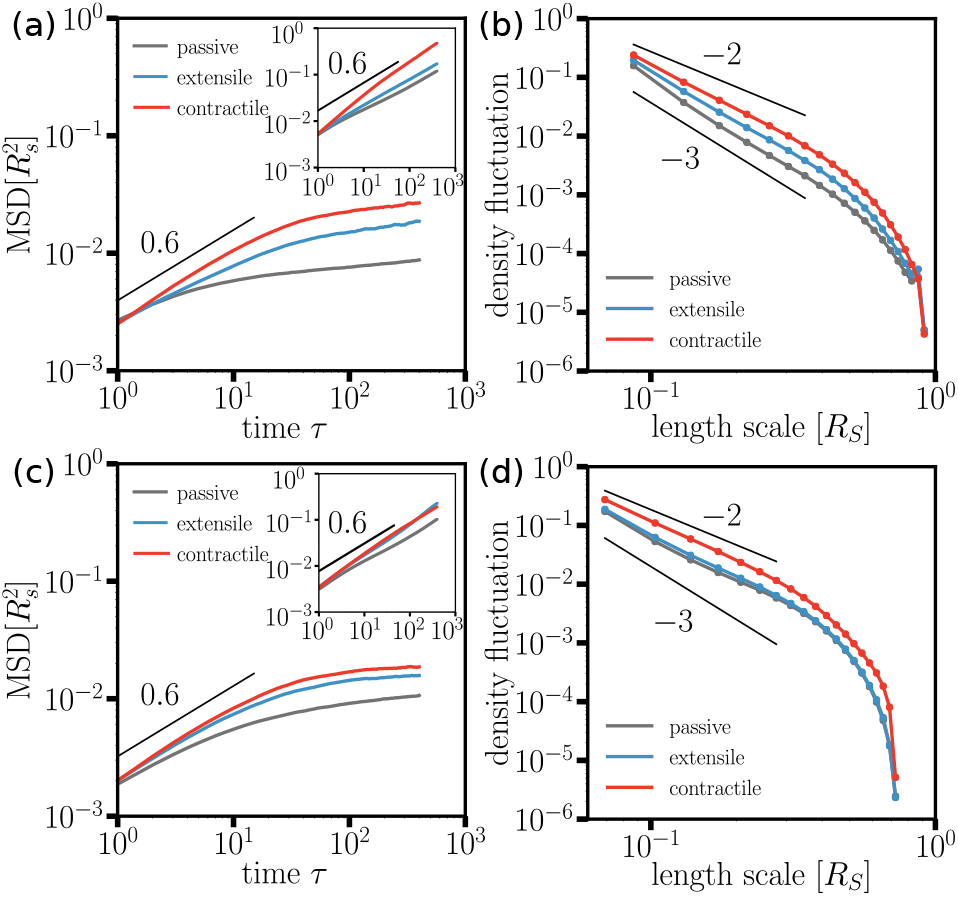
(a) MSD for the hard shell case with *N*_*C*_ = 2500, *N*_*L*_ = 50, and *M* = 5. For the inset, *N*_*C*_ = 0. (b) Density fluctuations for the same parameters as in (a). Figures (c) and (d) show the soft shell equivalent to (a) and (b).

Additionally, the contractile system exhibits larger displacements than the extensile system. We found that a broader spectrum of steady-state density fluctuations for the contractile system drive this behavior (Fig. 3 (b)). This generates regions of lower density into which the chain can move, leading to increased motility. The active cases exhibit anomalous density fluctuations, with the variance in the density falling off more slowly than inverse length cubed (in 3D). Finally, the MSD in the hard shell case is suppressed by more boundary bindings or crosslinks. For the soft shell case, we observe similar trends as the hard shell, except that the soft shell does not inhibit the potential gel-sol transition.

Next, we examine nuclear shape. In Figure 4, we plot the power spectrum of the shape fluctuations of the shell for a central cross-section as a function of wavenumber *q* for different motor strengths. We observe that the shape fluctuation spectrum is broad until saturating due to the discretization of the system. The decrease in the shape fluctuations is less significant for both the passive and extensile systems than for the contractile system with an approximate *q*^−2^ scaling, characteristic of membrane tension, for the former versus an approximate *q*^−3^ scaling for the latter. This difference could be due to the more anomalous density fluctuations in the contractile case, demonstrating that chromatin spatiotemporal dynamics directly impacts nuclear shape. We do not observe a *q*^−4^ contribution due to emergent bending, which was suggested by previous experiments [17] and simulations [23]. However, additional experiments measuring nuclear shape fluctuations of mouse embryonic fibroblasts (MEFs) also do not show a bending contribution (inset to Fig. 4 and see SM for materials and methods). Additionally, the amplitude of the shape fluctuations increases with motor strength, *N*_*C*_, and *N*_*L*_ (see SM).

**FIG. 4.**
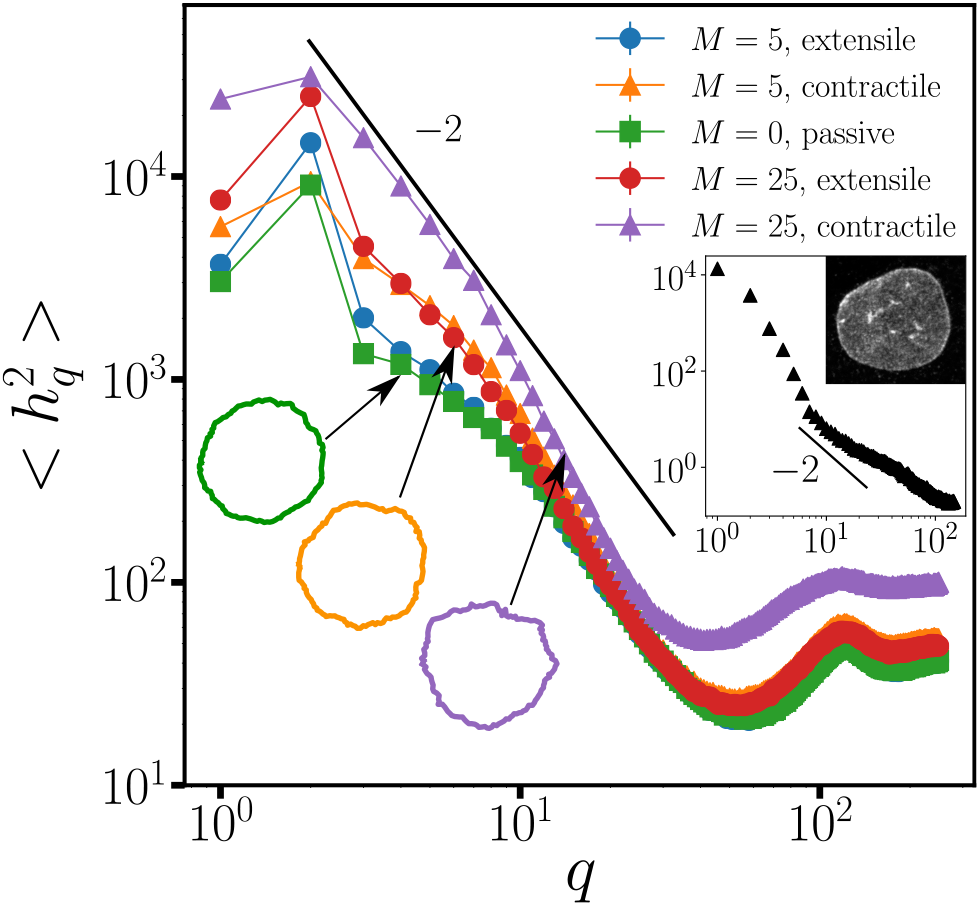
Power spectrum of the shape fluctuations with *N*_*L*_ = 50 and *N*_*C*_ = 2500 for the passive and both active cases. Different motor strengths are shown. The insets shows experimental data from mouse embryonic fibroblasts with an image of a nucleus with lamin A/C stained.

## Discussion

We have studied a composite chromatin-lamina system in the presence of activity, crosslinking, and number of linkages between chromatin and the lamina. Our model captures the correlated chromatin motion on the scale of the nucleus in the presence of both activity and crosslinks (Fig. 2). The deformability of the shell also plays a role. We find that global translations of the composite soft shell system contribute to the correlations. We observe anomalous diffusion for the chromatin (Figs. 3 (a) and (c)), as has been observed experimentally [22], with a crossover to a smaller anomalous exponent driven by the crosslinking [32]. Interestingly, the contractile system exhibits a larger MSD than the extensile one, which is potentially related to the more anomalous density fluctuations in the contractile case (Figs. 3 and (d)). Finally, nuclear shape fluctuations depend on motor strength and on amounts of crosslinking and chromatin-lamina linkages (Fig. 4). Notably, the contractile case exhibits more dramatic changes in the shape fluctuations as a function of wavenumber as compared to the extensile case.

Our short-ranged, overdamped model contrasts with an earlier confined, active Rouse chain interacting with a solvent via long-range hydrodynamics [4]. While both models generate correlated chromatin dynamics, with the earlier model, such correlations are generated only with extensile motors that drive local nematic ordering of the chromatin chain [4]. Moreover, our correlation lengths are significantly larger than those obtained in a confined active, heteropolymer simulation [15]. Activity in this earlier model is modeled as extra-strong thermal noise such that the correlation length decreases at longer time windows as compared to the passive case. This decrease contrasts with our results (Figs. 2 (d) and (h)) and experiments [3]. In addition, our model takes into account deformability of the shell and the chromatin-lamina linkages. Future experiments could potentially distinguish these mechanisms by looking for prominent features of our model, such as a dependence on chromatin bridging proteins and linkages to the lamina and effects of whole-nucleus motions.

Our modeling motivates further spatiotemporal studies of nuclear shape. Particularly interesting would be *in vivo* studies with vimentin-null cells, which have minimal mechanical coupling between the cytoskeleton and the nucleus. Vimentin is a cytoskeletal intermediate filament that forms a protective cage on the outside of the nucleus and helps regulate the nucleus-cytoplasm coupling and, thus, affects nuclear shape [8]. The amplitudes of the nuclear shape fluctuations in vimentin-null cells may increase due to a softer perinuclear shell or may decrease due to fewer linkages between the nucleus and the mechanically active cytoskeleton.

There are intriguing parallels between cell shape [34–36] and nuclear shape with cell shape being driven by an underlying cytoskeletal network—an active, filamentous system driven by polymerization/depolymerization, crosslinking, and motors, both individually and in clusters, that can remodel, bundle and even crosslink filaments. Given the emerging picture of chromatin motors acting collectively [26, 27], just as myosin motors do [37], the parallels are strengthened. Moreover, the more anomalous density fluctuations for the contractile motors as compared to the extensile motors could potentially be relevant in random actin-myosin systems typically exhibiting contractile behavior, even though either is allowed by a statistical symmetry [38]. On the other hand, distinct physical mechanisms may govern nuclear shape since the chromatin fiber is generally softer than cytoskeletal filaments and the lamina is stiffer than the cell membrane.

We now have a minimal chromatin-lamina model that can be augmented with additional factors, such as different types of motors—dipolar, quadrupolar, and even chiral, such as torque dipoles. Chiral motors may readily condense chromatin just as twirling a fork “condenses” spaghetti. Finally, it is now established that nuclear actin exists in the cell nucleus, yet its form is under investigation [39]. We propose that short, but stiff, actin filaments acting as stir bars can potentially increase the correlation length of micron-scale chromatin dynamics. Including such factors will help us further quantify nuclear dynamics to determine, for example, mechanisms for extreme nuclear shape deformations, such as nuclear blebs [40], and ultimately how nuclear spatiotemporal structure affects nuclear function.

JMS acknowledges financial support from NSF-DMR-1832002. JMS and AEP acknowledge financial support from a CUSE grant. EJB was supported by the NIH Center for 3D Structure and Physics of the Genome of the 4DN Consortium (U54DK107980), the NIH Physical Sciences-Oncology Center (U54CA193419), and NIH grant GM114190.

## Appendix A: Model

## 1. System and initialization

We use a Rouse chain with soft-core repulsion between each monomer capturing excluded volume effects to represent the chromatin. Since the chromatin is contained within the lamina, modeled as a polymeric shell, we present the protocol to obtain the initial configuration for the composite system. As shown in Fig. S1(left), we first implement a three-dimensional self-avoiding random walk in an FCC lattice for 5000 steps to generate the chain. We then surround the chain in a large polymeric, but hard, shell. To create the shell, we generate a Fibonacci sphere with 5000 nodes and identify 5000 identical monomers with these nodes. The springs between the shell monomers form a mesh and each shell monomer is connected to 4.5 other shell monomers on average. These monomers have same physical properties as the chain monomers in terms of size and spring strength.

We then shrink the shell (Fig. S1(center)) by moving the shell monomers inwards by the same amount. During the shrinking process, chain monomers interact with the shell monomers via the soft-core repulsion and, therefore, also move inwards. In addition, every chain monomer experiences thermal fluctuations and is constrained by elastic forces and soft-core repulsion forces. Once the shell radius reaches its destination radius after some time, we then thermalize the positions of the shell monomers and adjust rest length of springs respectively to make the mesh less lattice-like. We, thus, arrive at the initial configuration of the system Fig. S1(right). We obtain 100 such initialized samples to obtain an ensemble average for each measurement. The destination radius *R*_*s*_ is 10. We set the monomer radius to be *r*_*c*_ = 0.43089 so that the packing fraction *ϕ* is approximately 0.4 in the hard shell limit comparable to electron microscopy tomography experiments [1], simulations of chromatin confined within the nucleus [2], and theoretical estimates [3], while *ϕ* is smaller in soft-shell cases due to expansion as the shell monomers undergo thermal fluctuations.

## 2. Parameters

In our simulations, we use the set of parameters shown in Table 1.

**FIG. S1.**
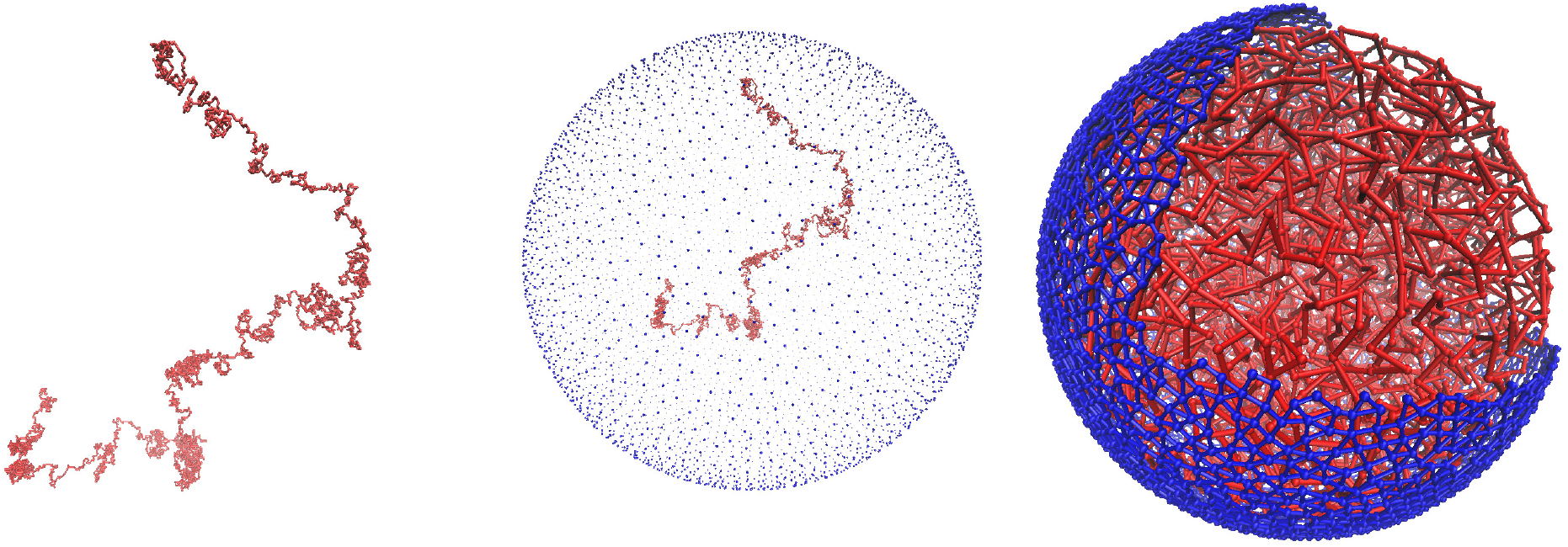
Left: The chain is initially generated via a self-avoiding random walk on an FCC lattice. Center: The chain is then enclosed in a Fibonacci sphere. Right: Composite system at time *τ* = 0.

|  |  |  |
| --- | --- | --- |
| Diffusion constant | $D$ | 1 |
| Thermal energy | $k_B T$ | 1 |
| Simulation timestep | $d\tau$ | $10^{-4}$ |
| Number of chain monomers | $N$ | 5000 |
| Radii of chain monomers | $r_c$ | 0.43089 |
| Number of shell monomers | $N_s$ | 5000 |
| Radii of shell monomers | $r_s$ | 0.43089 |
| Radius of hard shell | $R_s$ | 10 |
| Packing fraction | $\phi$ | 0.400 |
| Spring constant | $K$ | 140 |
| Soft-core repulsion strength | $K_{ex}$ | 140 |
| Number of motors | $N_m$ | 400 |
| Motor strength | $M$ | 5/25 |
| Turnover time for motors | $\tau_m$ | 0.05 |
| Number of crosslinks | $N_C$ | 0/100/500/1000/2000/2500 |
| Number of linkages | $N_L$ | 0/50/200/400/600 |
| Damping | $\xi$ | 1 |

We now address how the simulation parameters map to biological values. One simulation length unit corresponds to 1 *μ*m, one simulation time unit corresponds to 0.5 seconds, and one simulation energy scale corresponds to approximately 10^−21^ *J* = *k_B_T*, *T* = 300 K. With this mapping, the spring constant corresponds to approximately 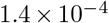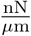 with a Young’s modulus for the chain of 0.28 Pa.

## Appendix B: Simulation results

## 1. Radius of gyration

For a polymer, the radius of gyration is defined as *R*_*g*_ = Σ(*r*_*i*_ − *r*_*cm*_)^2^/*N*, where *N* = 5000 is total chain monomer number. In the hard shell case, we fix the radius of the shell to *R*_*s*_ = 10. In the soft-shell case, the shell expands due to the thermal fluctuations and due to the activity of the chain inside. Fig S2 shows the radius of gyration of the chain (solid lines) and the average radius of shell (dashed lines) in the soft shell case as function of time. After a short-time initial expansion, both the chain’s and the shell’s respective radii reach a plateau by 100 *τ* for most parameters, indicating that the system is reaching steady state. Only for the zero crosslinks with contractile activity, does the radius of gyration continue to increase slightly over the duration of the simulation of 500 *τ*.

**FIG. S2.**
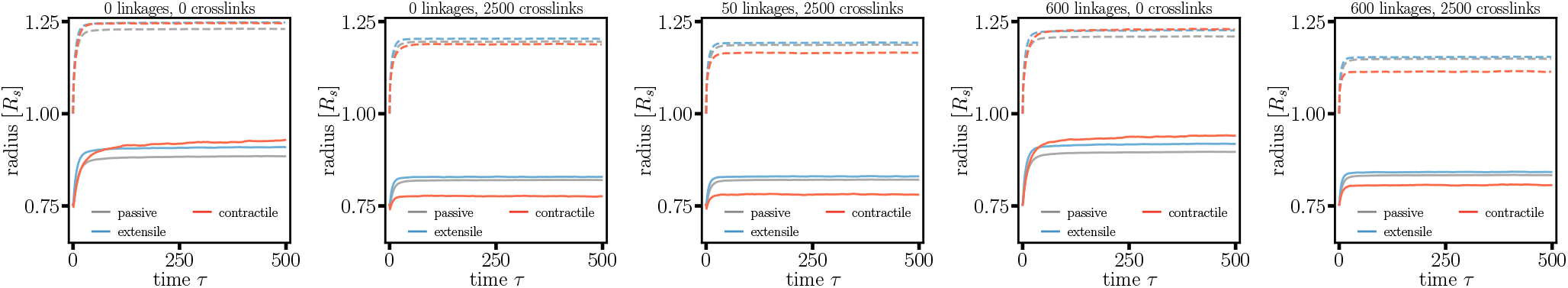
Radius of gyration of the chain (solid lines) and average radius of the shell(dashed lines) as function of simulation time for *N*_*C*_ = 2500 and *N*_*L*_ = 50 (middle figure). For contrast, *R*_*g*_ for *N*_*L*_ = 0, 600 and *N*_*c*_ = 0, 2500 are also plotted. Only the soft shell case are shown.

## 2. Self-contact probability

Since the globule radius is an averaged quantity, we also look for steady state signatures in the self-contact probability, which yields information about the chromatin spatial structure. More specifically, Hi-C allows one to quantify the local chromatin interaction domains at the megabase scale [4]. Such domains are stable across different eukaryotic cell types and species [5]. To quantify such interactions in the simulations, one determines the number of monomers in the vicinity of the *ith* chain monomer. In other words, one creates an adjacency matrix. This adjacency matrix is shown Fig. S3 for two examples. To compute the self-contact probability, one sets a threshold distance that a pair of monomers within that range is considered to be in contact. Then the fraction of contacted pairs for each polymeric distance 1, 2, 3, 4, … is calculated. This fraction as a function of polymeric distance is called the self-contact probability. See Fig. S4 for the self-contact probability for *N*_*L*_ = *N*_*C*_ = 0 at the beginning and at the end of the simulation for the soft shell case. While there is some change between the two, in Fig. S5, we show the self-contact probability for different times *τ* to demonstrate that after *τ* = 50, the probability does not change with time, implying a steady state.

**FIG. S3.**
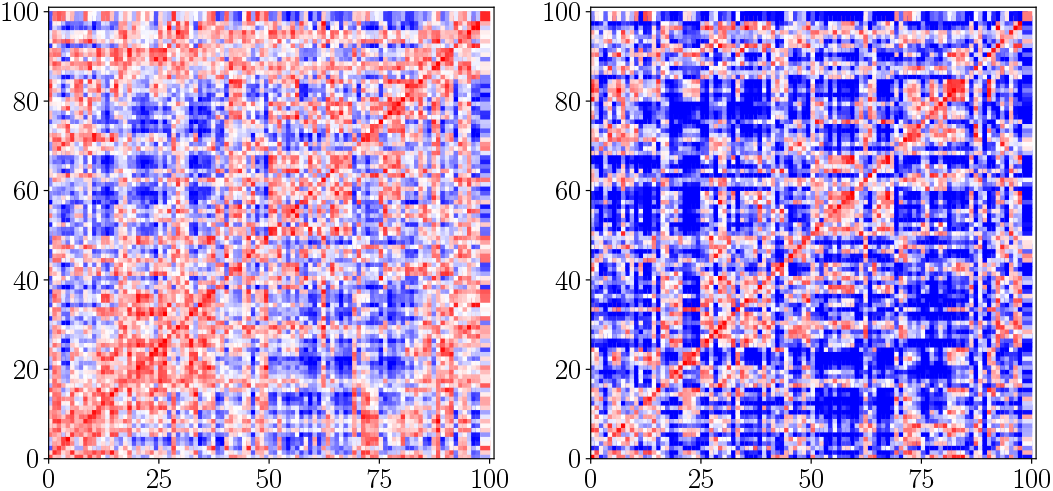
Contact map for a soft shell, contractile system with no linkages or crosslinks at the beginning and at the end of the simulation, i.e. *τ* = 0 and *τ* = 500.

## 3. Mean-Squared Displacement

To quantify the dynamics of the chain, we compute its mean-squared displacement (MSD) measured with respect to the center of mass of the shell. Fig. S6 plots the MSD of the chain during the duration of the simulation. At short time scales, the chain undergoes sub-diffusive motion and the MSD follows an exponent around *α* ≈ 0.6 for *N*_*C*_ = 2500 and *N*_*L*_ = 50. At longer time scales, the MSD crosses over to a smaller exponent. The value of the exponent depends on *N*_*C*_ and *N*_*L*_. In all cases, the active systems diffuse faster than the passive system, and contractile motors enhance diffusion more than extensile motors. The insets in Fig. S6 show the MSD for the center of mass of the chromatin chain for the soft shell. For the crosslinked, active chain, this MSD is slightly faster than diffusive.

**FIG. S4.**
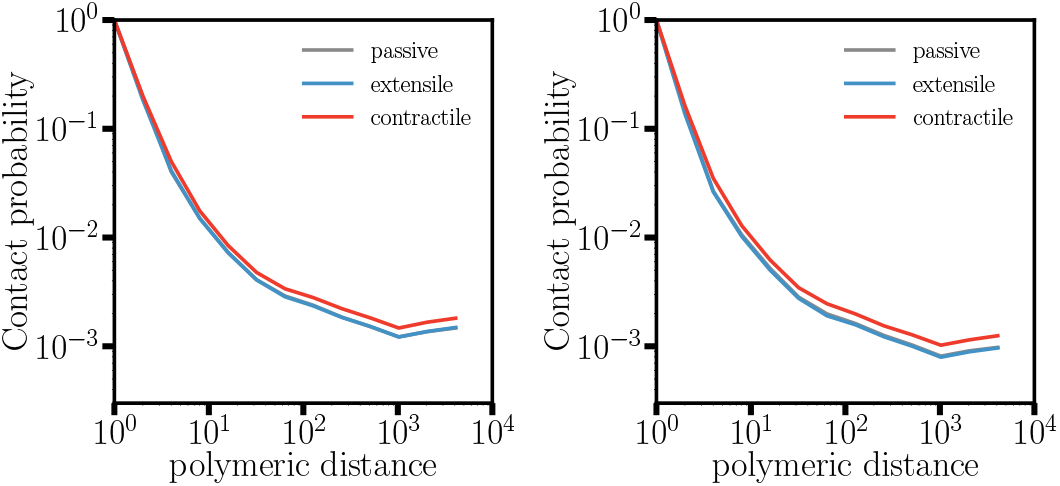
Self-contact probability for the hard shell case (left) and the soft shell case (right) with the latter corresponding to right figure in previous Fig. S3.

**FIG. S5.**
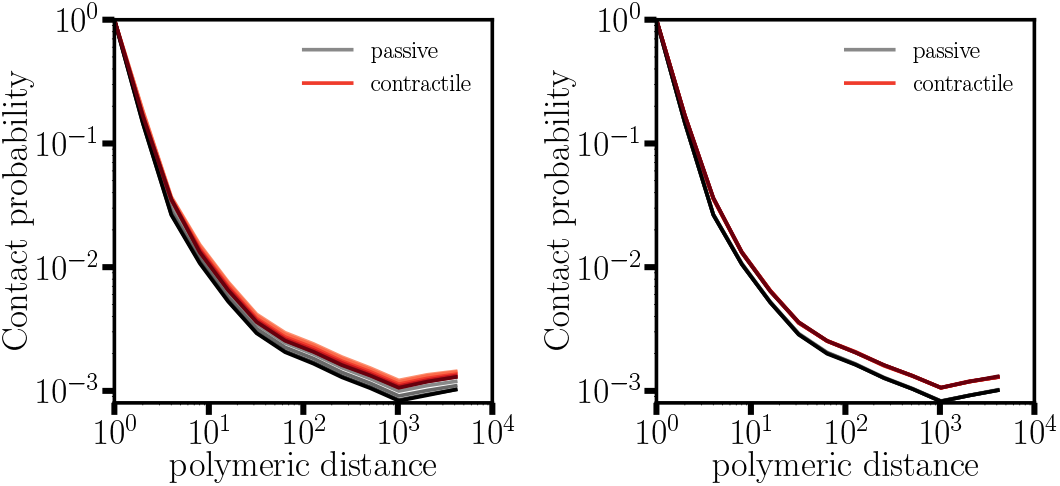
Self-contact probability at *τ* = 0, 1, 2, 10, 20, 50 *τ* (left) and for *τ* = 50 and *τ* = 300 (right) for soft shell passive and contractile systems with *N*_*C*_ = 2500 and *N*_*L*_ = 50.

## 4. Density fluctuations

The density fluctuations are computed in the following way:

- Select a spherical region in the system with radius *r*_*d*_ and count the number of monomers in that region.
- Randomly select spherical regions at other places with the same radius and count the monomers included.
- Compute the variance of counted monomer amount *σ*^2^ for this radius *r*_*d*_.
- Vary *r*_*d*_ and repeat the above three steps and obtain the variance for each *r*_*d*_.

We plot *σ*^2^ as a function of *r*_*d*_. Typically, for a group of randomly distributed monomers in three dimensions, the density fluctuations scale as 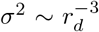. From Fig. S7 we see that the overall density fluctuations are broader in the active cases, as compared to the passive cases. Contractile motors induce more anomalous density fluctuations, particularly in the soft shell case.

## 5. Correlation function and correlation length

To evaluate the spatial and temporal correlation motion along the chain, we compute the spatial autocorrelation function. Suppose *d* 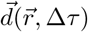 is the displacement of monomer at 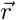 over time, 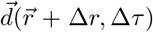 is the displacement of another monomer, which is located a distance Δ*r* away and over the same time window. We then use the function below to compute the correlation function:

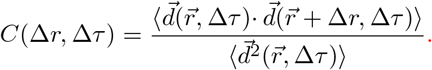

**FIG. S6.**
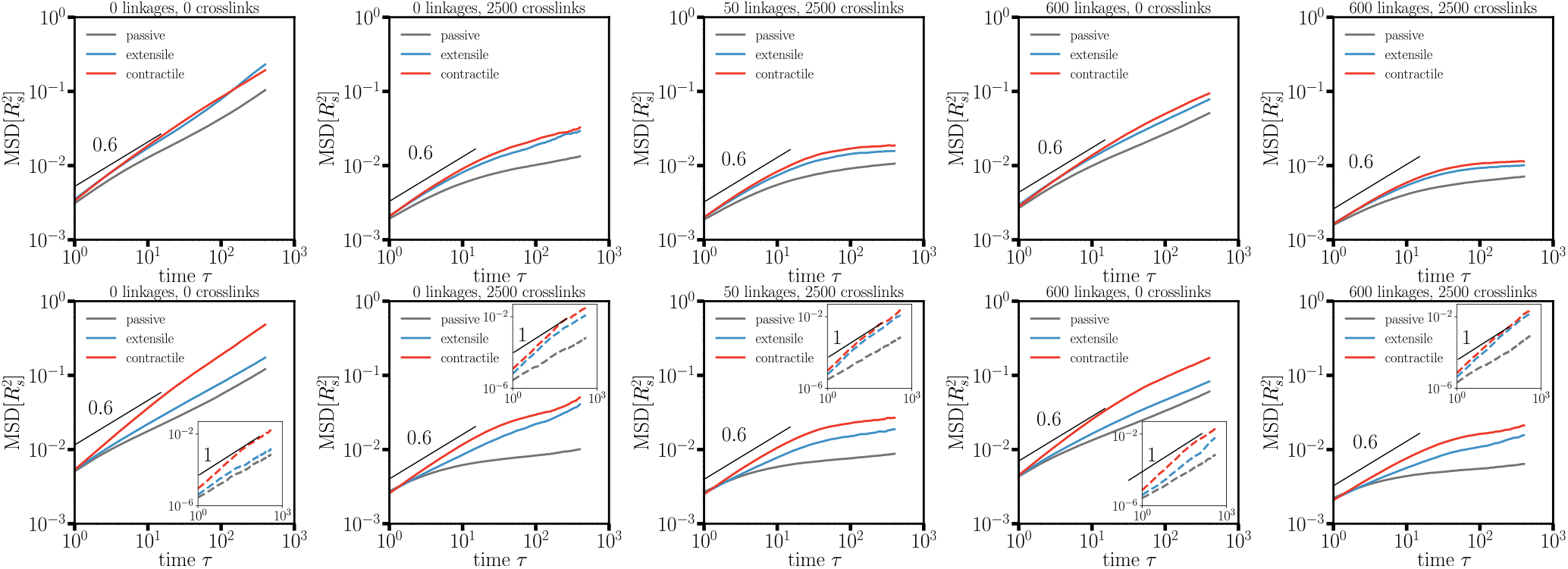
MSD as a function of time for *N*_*C*_ = 2500, *N*_*L*_ = 50, and *M* = 5 (middle column) and for four extreme cases (0 or 600 linkages, 0 or 2500 crosslinks) in the hard shell (top row) and the soft shell (bottom row). Insets are MSD plots of the center of mass of the chain.

From Ref. [6] we assume the correlation function follows 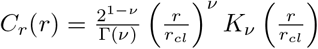, where *r*_*cl*_ is the extracted correlation length, *K*_*ν*_ is the Bessel of the second type of order *ν*, and *ν* is a smoothness parameter. Larger *ν* denotes that the underlying spatial process is smooth, not rough, in space. In Fig. S8 we show the correlated function computed from numerical simulations (dots) and the fitted correlation function from the above formula (lines) for different parameters. Lines from light to dark represent time windows from short to long (1 *τ*, 2 *τ*, 5 *τ*, 10 *τ*, 20 *τ*, 50 *τ*, 100 *τ*, 200 *τ*). We see that the numerical results with shorter time windows fit the formula better.

In Fig. S9, we plot the correlation length a function of linkage number *N*_*L*_ and crosslink number *N*_*C*_ over the short time window 5 *τ* and the long time window 50 *τ*. We observe that active motors clearly enhance the correlation length. It is also clear that presence of crosslinks also enhance correlation length. The correlation length is larger for the soft shell case. In the soft shell case, without subtracting the diffusion of the center of mass, the correlation length for the long time window spans almost the radius of the system. We note that the correlation length is reduced if we subtract the center of mass shell motion, however, it still remains larger than the hard shell case. A quasi-two-dimensional correlation length is computed from a slab-like region and is also shown for potential comparison to experimental results since, in the experiments, the correlated length is extracted using this method. There is not much difference between the three-dimensional correlation length and the two-dimensional correlation length with the center of mass of the shell subtracted. We also show the corrrelation length as a function of shell stiffness (with the COM of shell subtracted) to demonstrate the direct effect of shell stiffness on the correlated chromatin motion (see Fig. S10).

**FIG. S7.**
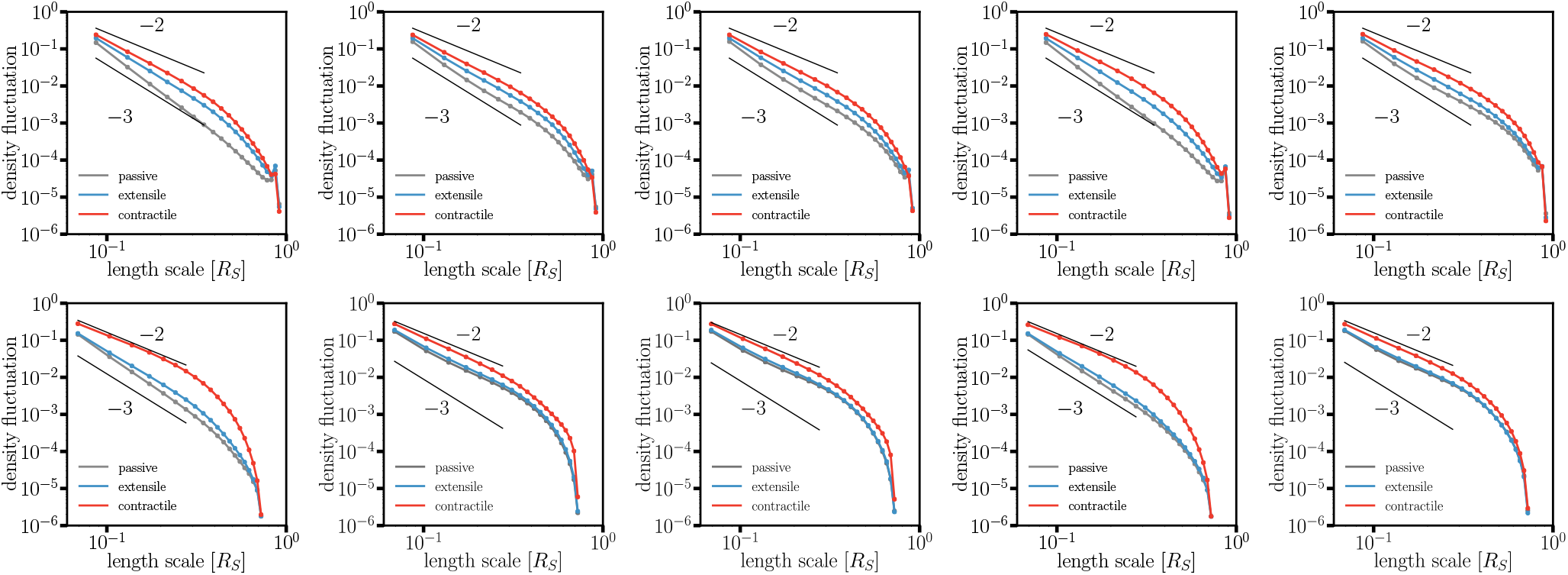
Density fluctuations for *N*_*C*_ = 2500 and *N*_*L*_ = 50 (middle column) and four extreme cases (0 or 600 linkages, 0 or 2500 crosslinks) in the hard shell (top row) and in the soft shell (bottom row). The arrangement of parameters is the same as in the previous figure.

**FIG. S8.**
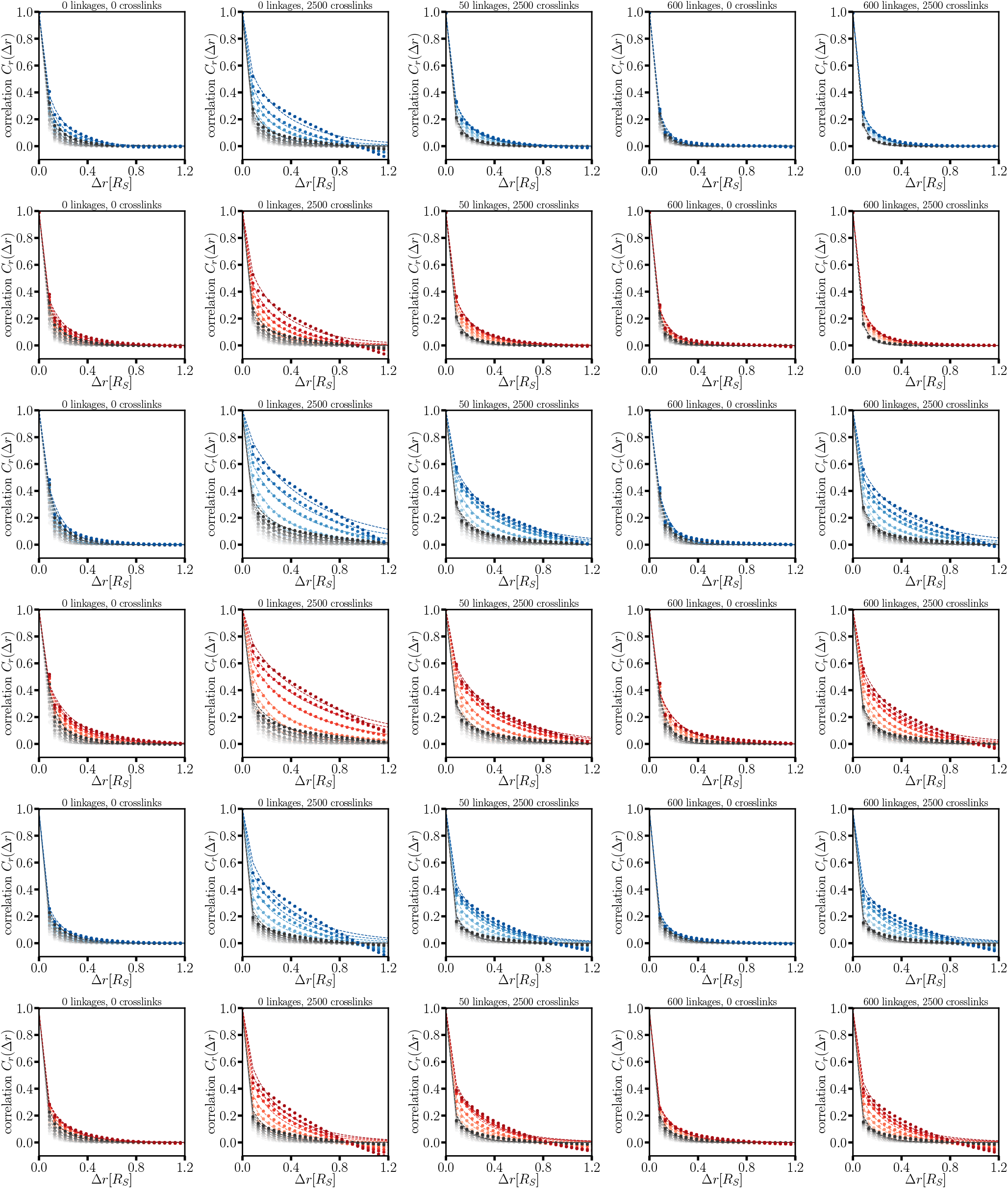
Correlation functions for *N*_*C*_ = 2500 and *N*_*L*_ = 50 (middle column) and four extreme cases (left column: 0 linkages and 0 crosslinks; second from left column: 0 linkages and 2500 crosslinks; second from right column: 600 linkages and 0 crosslinks; right column: 600 linkages and 2500 crosslinks). Top two rows: The three-dimensional correlation function for the hard shell; Middle two rows: The three-dimensional correlation functions for the soft shell; Bottom two rows: Two-dimensional correlation functions for the soft shell. Color varies from light to dark as time lag equals 1 *τ*, 2 *τ*, 5 *τ*, 10 *τ*, 20 *τ*, 50 *τ*, 100 *τ*, 200*, τ*, respectively. Symbols denote the numerical results, while the dashed line represent the fitted correlation functios. Greyscale: passive. Bluescale: active with extensile motors. Redscale: active with contractile motors.

**FIG. S9.**
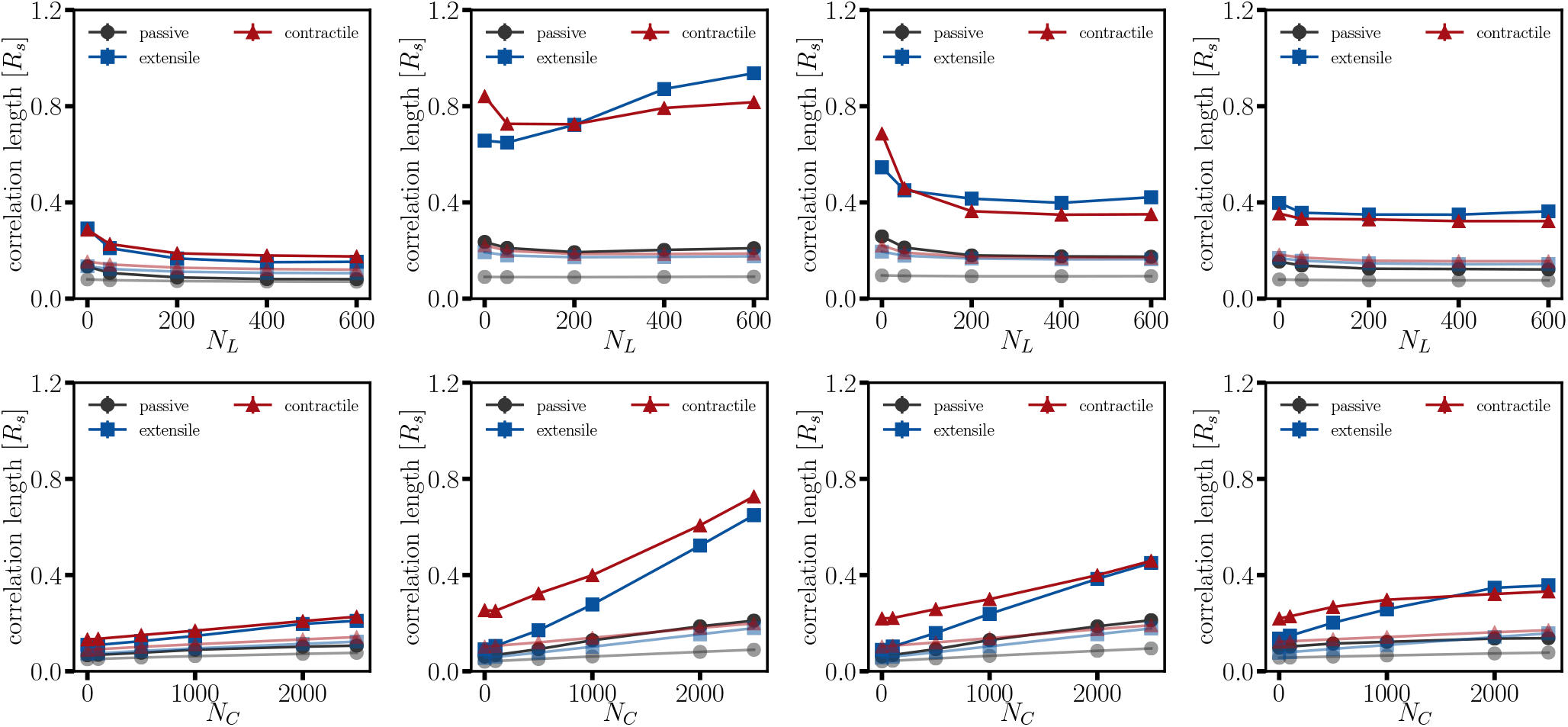
Plot of correlation length as function of linkage number *N*_*L*_ (top row) or crosslink number *N*_*C*_ (bottom row) for time windows 5 *τ* (light) and 50 *τ* (dark). From left to right columns: The three-dimensional correlation length for the hard shell; the three-dimensional correlation length for the soft shell; three-dimensional correlation length for the soft shell with the COM motion subtracted; two-dimensional correlation length for the soft shell with the COM motion subtracted.

**FIG. S10.**
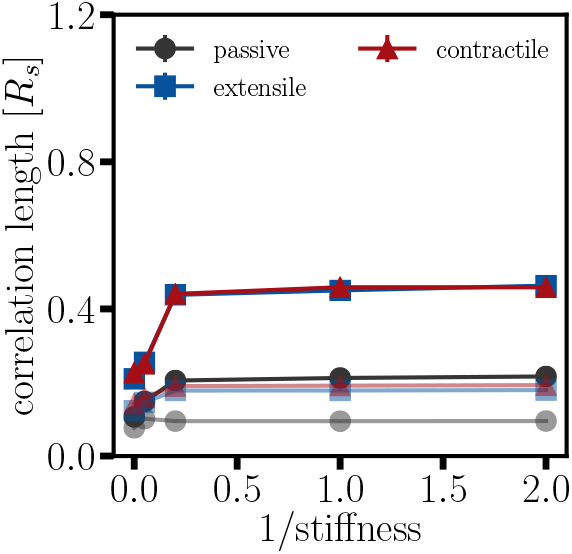
Plot of the correlation length as a function of shell stiffness for time windows Δ*τ* = 5, 50. Here *N*_*C*_ = 2500, *N*_*L*_ = 50, and *M* = 5.

## 6. Shape fluctuations

To evaluate shape fluctuations of the shell, we compute it in two ways. First, in order to compare with experimental measurements, we select a random slab through the center and project the coordinates of the shell monomers in the slab to the plane where slab lies. Then, we compute the fast-Fourier-transform (FFT) for spatial deviations of these monomers from the average radius with the deviations with *h*_*q*_ denoting the Fourier transform of the deviation with respect to wavenumber *q*. In Fig. S11, the power spectrum of the shape fluctuations for the passive and extensile cases follow a decay exponent of 2, as expected for a stretchable shell [7]. The spectrum of the shape fluctuations increases monotonically with the number of crosslinks. The specturm varies more dramatically with contractile motors as compared to extensile motors. Moreover, the shape fluctuation spectrum also eventually saturates as a function of chromatin-lamina linkage number. In S12 we compute the spectrum of the shape fluctuations as characterized by the spherical harmonic functions (the *Y_lm_*s with *l* as the dimensionless spherical wavenumber). We obtain similar trends as in Fig. S11. Finally, in Fig. S13, we plot the spectrum for different motor strengths and different shell stiffnesses.

**FIG. S11.**
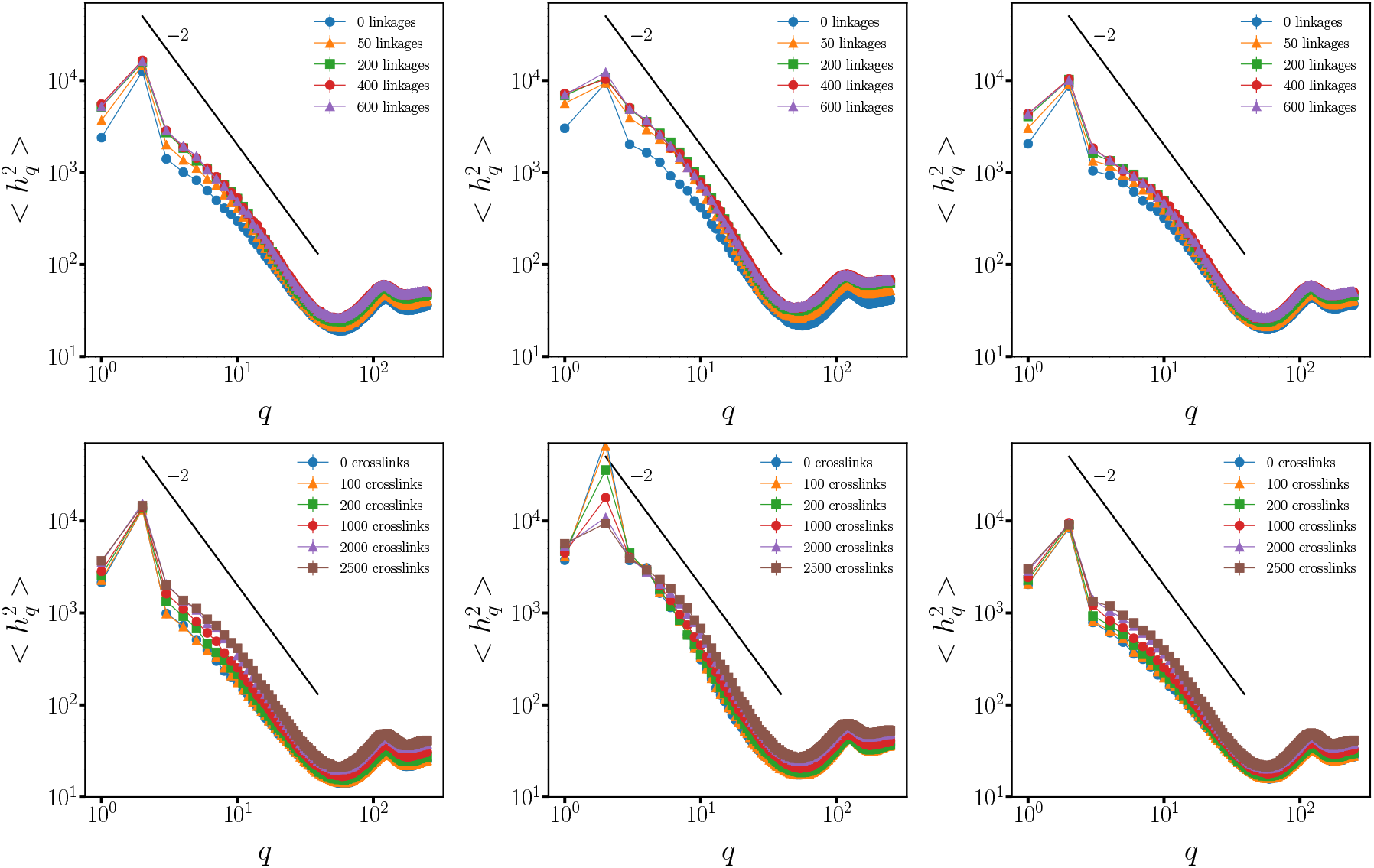
Power spectrum of the shape fluctuations of a random slab for different boundary linkages (top row) and crosslinks (bottom row). Left column: Extensile motor case. Middle column: Contractile motor case. Right column: Passive case.

## Appendix C: Experiments

To measure nuclear shape fluctuations in live cells, the wild-type mouse embryonic fibroblasts (mEFs) were kindly provided by J. Eriksson, Abo Akademi University, Turku, Finland. Cells were cultured in DMEM with 25 mM Hepes and sodium pyruvate supplemented with 10% FBS, 1% penicillin/streptomycin, and nonessential amino acids. The cell cultures were maintained at 37 degrees C and 5% CO_2_.

Cell nuclei were fluorescently labeled by transient transfection with pEGFP-C1-NLS, 48 h before imaging. Cell nuclei were imaged at 2-min increments for 2 h by using wide-field fluorescence with a 40× objective. To quantify the structural features of nuclei, we traced the contour, *r*(*θ*), of the NLS-GFP labeled nuclei at each time point. The shape of the nucleus was identified using a custom-written Python script, and its contour was interpolated from 0 to 2*π* by 150 points. Next, the shape fluctuations were calculated as *h*(*θ*) = *r*(*θ*) − *r*_0_, where *r*_0_ is the average radius for each cell at each time point. The wave number-dependent Fourier modes of the fluctuations, *h*_*q*_, were obtained as Fourier transformation coefficients, as described in Ref [8].

The shape fluctuations were quantified for each cell by computing the Fourier mode magnitude square *h*^2^(*q*) and averaging over each time point. The average shape fluctuations as shown in Fig. 4 in the main text was taken as the average over 15 cells per condition from two independent experiments.

**FIG. S12.**
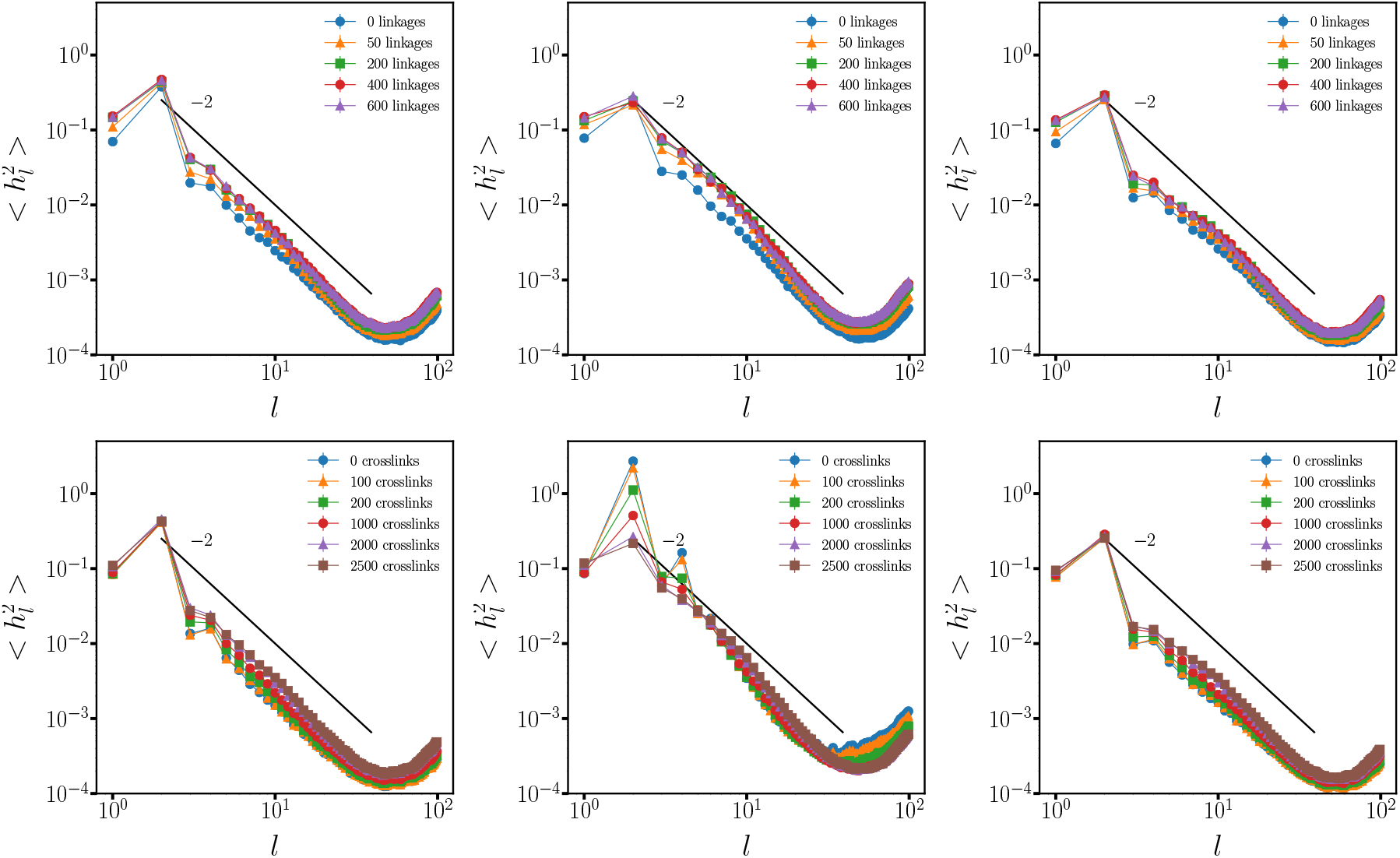
Power spectrum of the shape fluctuations in spherical harmonics, where *l* is the dimensionless spherical wavenumber for different chromatin-lamina linkages (top row) or crosslinks (bottom row). Left column: Extensile motor case. Middle column: Contractile motor case. Right column: Passive case.

**FIG. S13.**
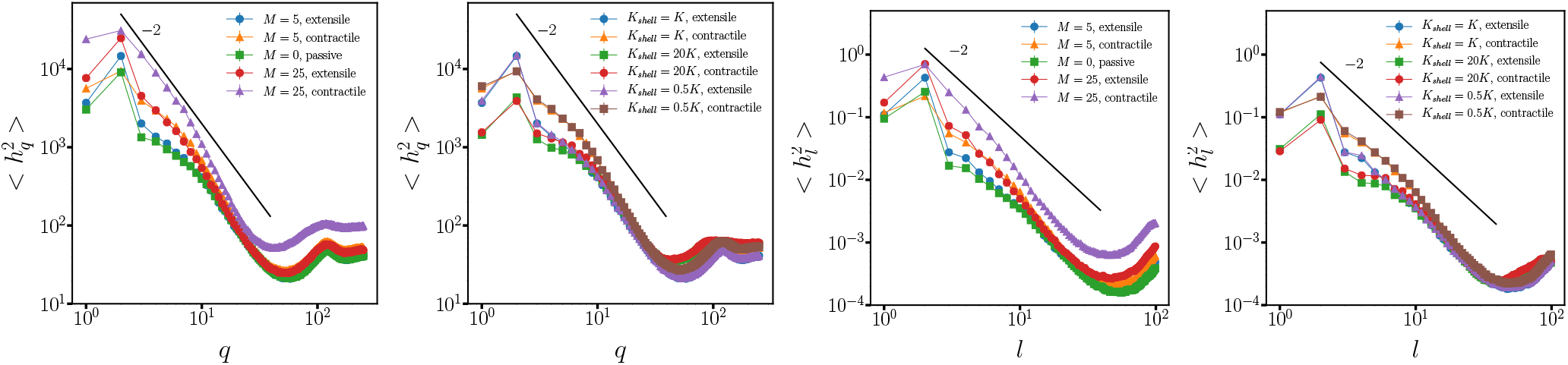
Power spectrum of the shape fluctuations for different motor strengths or shell stiffnesses. Left two: *q* plot. Right two: *Y*_*lm*_ plot.

